# Seasonal Abundance and Spatial Pattern of Distribution of *Liriomyza trifolii* (Diptera: Agromyzidae) and Its Parasitoid on Bean and Squash in South Florida

**DOI:** 10.1101/2021.02.11.430801

**Authors:** Shashan Devkota, Dakshina R. Seal

## Abstract

American serpentine leafminer, *Liriomyza trifolii*, is a polyphagous insect pest that feeds on a wide range of vegetable and ornamental plants around the world. To develop an effective IPM program, information on the seasonal field distribution and population dynamics of leafminer and its parasitoids is very important. Therefore, seasonal abundances and spatial distributions of, *L. trifolii* on snap bean and squash were studied during four crop growing periods between 2013 to 2015 in Homestead, Florida. The mean numbers of mines, larvae, pupae, emerged adults, and parasitoids on snap bean were highest at 2 weeks after planting during all four growing periods. Whereas, the mean numbers of mines, larvae, pupae, emerged adults, and parasitoids on squash were highest at 3 weeks after planting during all four growing periods. *L. trifolii* distributions tended to be aggregated (1 < *b/β*) on snap bean at 2 weeks after planting during all four growing periods but had uniform (1 > *b/β*) distributions on squash at 2 weeks after planting during all four growing periods. Similar results were seen on the distribution of leafminer parasitoids on both bean and squash.

---

American serpentine leafminer, *Liriomyza trifolii*, is a polyphagous insect pest that feeds on a wide range of vegetable and ornamental plants around the world (Seal et al. 2006, Parrella 1987, Spencer, 1973). Damage is caused by adult females and larvae. Adult females make several punctures in the leaf, using their ovipositor. The punctures are made for feeding and egg laying. Major damage to the plant is caused by larval feeding. Leafminer larvae feeds on the mesophyll layer of leaves, which reduces the photosynthetic area. *L. trifolii* completes development from egg to adult in 19 days at 25°C (Leibee 1984). Because of a short development time, leafminer can produce multiple generations per season.

Biological control often plays a major role in managing *L. trifolii* (Burgess), which hosts more than 40 parasitoid species (Waterhouse and Norris 1987, Patel et al. 2003). In vegetable and ornamental production systems, *L. trifolii* is often considered a secondary pest, but its status has been raised to primary pest because excessive use of pesticides have reduced the natural enemies that usually regulate its population. If natural enemy species are sufficiently abundant, they can limit herbivore populations, which can allow plant communities to grow until they are limited by competition (Rosenheim et al. 1993, Colfer and Rosenheim 1995, Sher et al. 2000). Hence, information on parasitoid density and composition throughout the year, and their effects on leafminer density can help in developing an IPM program.

Changes in environmental factors, both biotic and abiotic, over time strongly affect leafminer development (Leibee 1985). For example, a female leafminer fly may lay up to 300 eggs per lifetime, which is over a span of 17 days at 25 °C (Charlton & Allen 1981). This rapid egg production may facilitate population increase (Jong and Rademaker 1991). Weather conditions including temperature, humidity, precipitation, and wind have are some of the most important causes of dramatic changes in pest abundance in an ecosystem (Risch 1987, Nestel et al. 1994). Changes in weather parameters may directly influence the physiology and behavior (locomotion and dispersal) of an insect and indirectly affect the insect population because of changes in the host plants and the behavior of its natural enemies (Martinat 1987, Nestel et al. 1994). In Lebanon, leafminer populations were reported to be reduced because of high mean temperatures in September and October (Hammad and Nemer 2000). Alternatively, Li et al. (2011) recorded increased leafminer populations in December and January, when mean temperatures were relatively low (21-23°C). Rainfall and humidity may also affect leafminer population. For instance, Shepard et al. (1998) found that leafminer populations on potato were relatively low during the dry season.

Cultivated crops are principal reproductive and feeding hosts of leafminer. However, when cultivated hosts are absent, leafminer tend to invade alternate weed hosts typically found nearby in the fields and returns to the main crops after they are re-planted. Knowledge of crop biology and ecology is also important in developing an IPM program for managing leafminer population. Therefore, to develop an effective IPM program, information on the seasonal field distribution and population dynamics of leafminer and its parasitoids is very important. The specific objectives of this study were: (a) to determine seasonal abundance of leafminer and its parasitoids on bean and squash in four plantings; (b) to determine their spatial distribution in four plantings.

## Materials and Methods

All the studies were conducted at two sites separated by 1 Km within the Tropical Research and Education Center, Homestead, FL. Snap bean (*Phaseolus vulgaris* L. ‘Prevail’) was planted at site 1 and squash (*Cucurbita pepo* L. ‘Enterprise’) was planted at site 2. Each crop was planted on four dates at the respective sites: Oct 26 (Oct-Nov 2013), May 10 (May-June 2014), Sep 6 (Sep-Oct 2014), and Nov 28 (Nov 2014-Jan 2015). Snap bean and squash seeds (Syngenta Seeds Inc., Othello, WA) were planted in a 92 m x 10 m field comprised of 6 raised beds each measuring 92 m x 1 m. Centers of adjacent beds were separated by 0.91 m. Each bed was divided into eight 11.5 m plots; hence, there were 48 plots. Snap bean and squash seeds were directly seeded on raised beds (1 m wide, 0.15 m high) covered with 1.5 ml thick black-and-white polyethylene mulch (Grower’s Solution Co., Cookeville, TN). 3-5 seeds were sown in a hole, 1.5 cm deep. Planting holes were spaced 25 cm within the row and 1 m between adjacent rows. A pre-plant herbicide, Halosulfuron methyl (Sandea^®^, Gowan Company LLC., Yuma, Arizona) was applied at 51.9 g / ha 21 days before planting to control weed emergence. Crops were fertilized applying granular fertilizer 6:12:12 (N: P: K) at 1345 kg/ha in a 10 cm-wide band on both sides of the raised bed center and was incorporated before placement of plastic mulch. Additionally, liquid fertilizer 4: 0: 8 (N: P: K) was also applied at 0.56 kg N / ha / day through a drip system at 3, 4, and 5 weeks after planting. Plants were irrigated every day for one hour to deliver water (1.25 cm) using two parallel lines of drip tube (T-systems, DripWorks, Inc., Willits, California), spaced 30 cm apart and parallel to the bed center, having an opening at every 13 cm. The fungicides, Chlorothalonil (Bravo^®^, Syngenta Crop Protection, Inc., Greensboro, NC) at 1.75 liter/ha and Copper hydroxide (Kocide^®^ 3000, BASF Ag Products, Research Triangle Park, NC) at 0.8 l/ha were sprayed every two weeks, using 655 l/ha at 207 kpa, to prevent fungal diseases. To control melonworms and pickleworms in squash, *Bacillus thuringiensis* based insecticides, Dipel DF^®^ (var. kurstaki) at 1.1 Kg/ha and Xentari DF^®^ (*B. thuringiensis* var. kurstaki) at 1.2 liters/ha (Valent Biosciences Corporation, Libertyville, IL), were used in weekly rotation.

### Seasonal Abundance of Leafminer Mines, Larvae, Adults, and Parasitoids

Seasonal abundance of leafminers was studied using snap beans and squash in four plantings. Sampling began 15 days after planting, when bean plants had two primary leaves fully unfolded. Five plants from each plot were randomly selected and one full grown leaf from the bottom stratum of each plant was sampled. Thus, five leaves were collected from each plot. All leaves from a plot were placed in a plastic pot (10 cm diameter and 15 cm depth) which was marked with date and plot number. The samples were then transported to the IPM laboratory and checked under a binocular microscope at 10X to record numbers of mines and larvae per leaf. Leaves were returned to the same pot and placed at room environment 25 ± 5 °C, 75 ±5% RH, and 14 h: 10 h (L: D) for further studies. All samples were checked at 24 h intervals for larvae and pupae and continued until the emergence of the last pupae. Pupae from each samples were placed separately in a Petri dish (10 cm diameter) marked with date and plot number and lined at the bottom with a moisten filter paper to prevent desiccation. Petri dishes with pupae were observed daily for adult and parasitoid emergence. Numbers of adult leafminer flies and parasitoids were recorded by date and plot number.

### Statistical analyses

Seasonal abundance data were analyzed independently for each planting by oneway analyses of variance (ANOVAs) using PROC MIXED in the SAS System (PROC MIXED, SAS Institute 2013). This system provides a very flexible modelling environment for handling a variety of problems involved with using subjects repeatedly. To normalize the error variances, all data were square-root transformed (⎷x+0.25) before the analyses. Repeated measures ANOVAs were used (PROC MIXED) because the same multiple treatments were surveyed on different dates. ANOVAs comparing mean numbers of mines, larvae, pupae, adults, and parasitoids were followed by Tukey-Kramer procedures for mean separation (*P* < 0.05) (SAS Institute 2013).

### Spatial Distribution

Spatial distribution for *L. trifolii* and its parasitoids were studied in the same field where abundance was studied. The data collected for abundance were also used for determining spatial distribution of *L. trifolii* and its parasitoid. In the present study, I used three different plot sizes to compare distribution pattern. The plot sizes were: 1) 23 m^2^ plots which were the combination of two adjacent initial sections; 2) 46 m^2^ plots which were the combination of four adjacent initial sections; and 3) 92 m^2^ plots which were the combinations of eight adjacent initial sections. Accordingly, data were pooled from 2, 4 and 8 initial sections, respectively.

### Statistical analysis

Spatial distribution was determined by using Taylor’s power law given by Equation 1 (Taylor 1961) and Iwao’s patchiness regression given by Equation 2 (Iwao 1968).

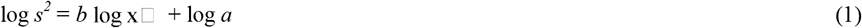

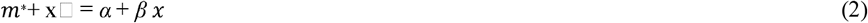

In Equation 1, Slope (*b*) is the index of aggregation, s^2^ is the variance, x□ is the mean number of leafminers, and *a* is the factor related to variability of sample size.

In Equation 2, m* is mean crowding index given by Lloyd (1967) which is ratio of sample variance (s^2^) and mean (x□). The slope *β*, which is similar to *b* value in Taylor’s power law, is the density of the contagiousness coefficient. *α* (intercept) is an index of basic contagion or tendency of insects towards crowding.

Both *b* and *β* in Taylor’s power law and Iowa’s patchiness regression, respectively, are indices of aggregation. Aggregate distributions resulted when *b* or *β* were significantly greater than 1, random when *b* and *β* were not significantly different from 1, and uniform (regular) when *b* and *β* values were significantly less than 1. The significance of slope *b* and β was determined by using student t-tests. Estimation of regression patterns were done by PROC GLM (SAS Institute Inc. 2013). Evaluation of the goodness of fit of the data for each linear model was done by an r^2^ value.

## Results

Seasonal Abundance of Leafminer Mines, Larvae, Adults, and Parasitoids

### Planting 1 (October - November 2013)

#### Site 1 (bean)

In Site 1 of Planting 1, the mining activity of *L. trifolii* on bean was significantly affected by sampling date (*F*_3, 141_ = 187.53, *P* < 0.0001) (Figure 1). The mean numbers of mines (53.33 ± 2.32 mines / 5 leaves) at 2 weeks after planting (Nov 9) were significantly higher than at 3 weeks (Nov 16), 4 weeks (Nov 23), or 5 weeks (Nov 30) after planting. The mean numbers of mines were the lowest (8.58 ± 0.46 mines / 5 leaves) when plants were 5 week old.

**Figure 1.**
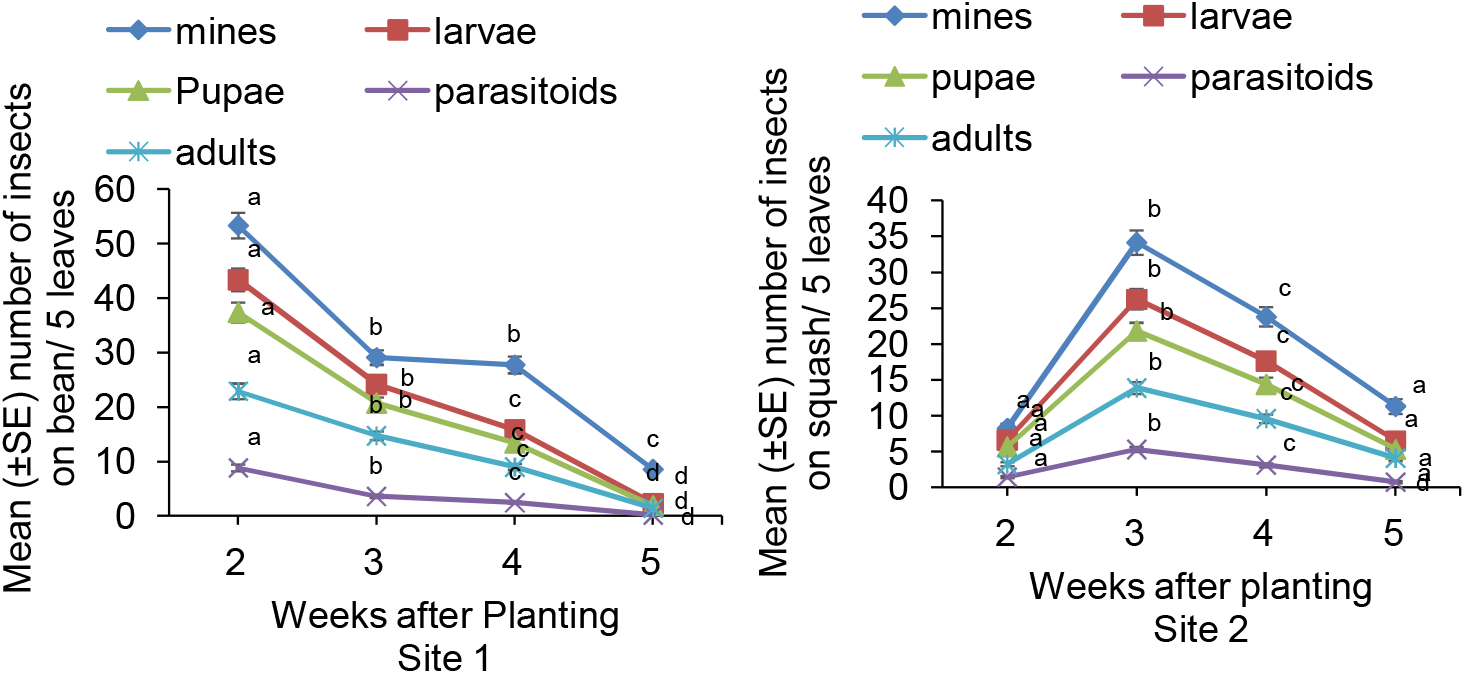
Planting 1 (26 Oct to 30 Nov 2013), Site 1 (bean) and Site 2 (squash) abundance of *L. trifolii* and its parasitoids (mean ± SE / 5 leaves). Means with same letters across the sample dates do not differ significantly based on a Tukey-Kramer test (*P* > 0.05).

Similarly, the mean numbers of larvae, pupae, emerged adults, and parasitoids were significantly affected by sampling dates (Larvae: *F*_3, 141_ = 288.63, *P* < 0.0001; Pupa: *F*_3, 141_ = 280.83, *P* < 0.0001; Adults: *F*_3, 141_ = 151.59, *P* < 0.0001; and parasitoids: *F*_3, 141_ = 191.99, *P* < 0.0001, respectively) (Figure 1). The mean numbers of larvae, pupae, emerged adults, and parasitoids were highest at 2 weeks (43.37 ± 2.08, 37.37 ± 1.83, 22.89 ± 1.42, and 8.85 ± 0.59 / 5 leaves, respectively) and lowest at 5 weeks after planting (2.31 ± 0.26, 1.91 ± 0.22, 1.43 ± 0.18, and 0.20 ± 0.24). The parasitoids recorded from leaf samples at site 1 were *Opius dissitus, Diglyphus sp., Euopius* sp. and *Diaulinopsis callichroma. O. dissitus* was the most abundant parasitoid and was about 70% of total population.

#### Site 2 (squash)

In Site 2 of Planting 1, the mean numbers of mines on squash were significantly affected by sampling dates (*F*_3, 141_ = 81.13, *P* < 0.0001) (Figure 1). The mean numbers of mines on squash (8.22 ± 0.692 mines / 5 leaves) were significantly lower at 2 weeks after planting (Nov 9). The mean numbers of mines (34.18 ± 1.69 mines / 5 leaves) increased significantly and reached the peak at 3 weeks after planting (Nov 16). Relative to the third week, numbers of mines then dropped significantly by 4 weeks after planting (23.79 ± 1.357 mines / 5 leaves, Nov 23). Again, at 5 weeks relative to the fourth week, the numbers of mines (11.29 ± 1.035) decreased significantly (Figure 1).

Similarly, sample dates significantly affected mean numbers of larvae, pupae, emerged adults, and parasitoids (Larvae: *F*_3, 141_ = 85.74, *P* < 0.0001; Pupae: *F*_3, 141_ = 87.35, *P* < 0.0001; Adults: *F*_3, 141_ = 79.57, *P* < 0.0001; and Parasitoids: *F*_3, 141_ = 59.79, *P* < 0.0001) (Figure 1). Mean numbers of larvae, pupae, and parasitoids were highest at 3 weeks after planting. *Opius dissitus, Diglyphus sp., Euopius* sp. and *Diaulinopsis callichroma* were the parasitoids recorded from site 2. *O. dissitus* was the most abundant parasitoid and was about 60% of total population.

### Planting 2 (May-June 2014)

#### Site 1 (bean)

Activity of *L. trifolii* in Planting 2 was similar to Planting 1. The mean numbers of mines, larvae, pupae, emerged adults and parasitoids were significantly affected by sample dates (Mines: *F*_3, 141_ = 396.23, *P* < 0.0001; Larvae: *F*_3, 141_ = 516.41, *P* < 0.0001; Pupae: *F*_3, 141_ = 517.30, *P* < 0.0001; Adults: *F*_3, 141_ = 392.71, *P* < 0.0001; Parasitoids: *F*_3, 141_ = 258.48, *P* < 0.0001) (Figure 2). The mean numbers of mines, larvae, pupae, emerged adults, and parasitoids (80.20 ± 3.63, 71.77 ± 3.41, 66.58 ± 3.24, 45.60 ± 2.68, and 16.10 ± 1.08 / 5 leaves; respectively) were highest at 2 weeks after planting (May 24) (Figure 2). These numbers were the highest among all sample dates across all seasons. There was a steep drop in mean numbers of mines, larvae, pupae, emerged adults, and parasitoids (27.89 ± 1.42, 20.52 ± 1.06, 16.77 ± 0.82, 11.31 ± 0.67, and 3.16 ± 0.26 / 5 leaves; respectively) at 3 weeks after planting (May 31). The mean numbers of mines, larvae, pupae, adults and parasitoids gradually decreased and were the lowest (7.33 ± 0.63; 2.43 ± 0.28; 2.04 ± 0.22; 1.47 ± 0.20; and 0.27 ± 0.07 / 5 leaves; respectively) at 5 weeks after planting (June 14) (Figure 2). *Opius dissitus, Diglyphus sp.*, and *Diaulinopsis callichroma* were the parasitoids recorded at site 1 in planting 2. *O. dissitus* was the most abundant among all the parasitoids recorded.

**Figure 2.**
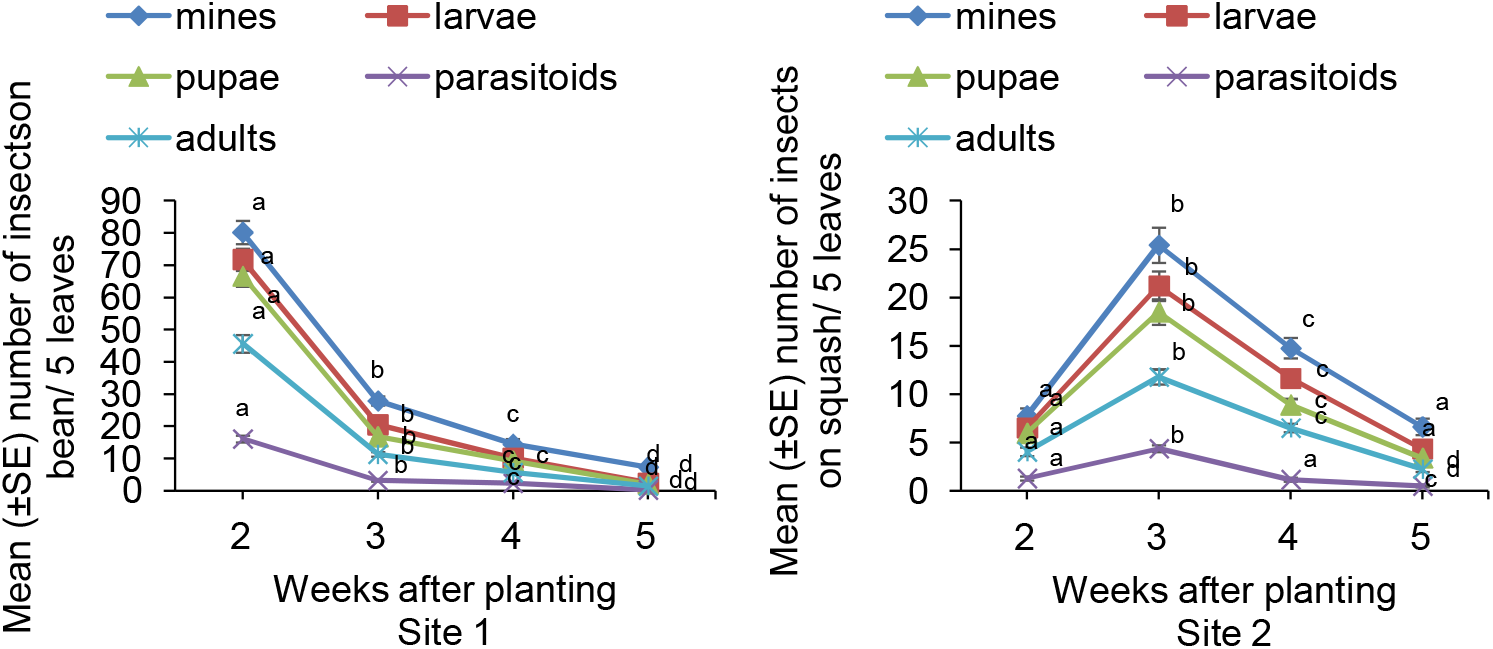
Planting 2 (10 May to 14 June 2014), Site 1 (bean) and Site 2 (squash) abundance of *L. trifolii* and its parasitoids (mean ± SE / 5 leaves). Means with same letters across the sample dates do not differ significantly based on a Tukey-Kramer test (*P* > 0.05).

#### Site 2 (squash)

Similar to Planting 1, the mean numbers of mines on squash, in Planting 2, were significantly affected by sample dates (*F*_3, 141_ = 52.76, *P* < 0.0001) (Figure 2). At 3 weeks after planting (May 31), the mean numbers of mines (25.41 ± 1.83 mines / 5 leaves) were significantly higher than at other sample dates (Figure 2). At 4 weeks after planting (June 7), the mean numbers of mines had dropped significantly (14.77 ± 1.04 mines / 5 leaves) compared to the third week, and at 5 weeks after planting (June 14), number of mines again dropped significantly (6.64 ± 0.84 mines / 5 leaves) compared to week 4 reaching lowest value. (Figure 2).

Similarly, sample date significantly affected mean numbers of larvae, pupae, emerged adults, and parasitoids (Larvae: *F*_3, 141_ = 65.04, *P* < 0.0001; Pupae: *F*_3, 141_ = 68.98, *P* < 0.0001; Adults: *F*_3, 141_ = 63.17, *P* < 0.0001; and Parasitoids: *F*_3, 141_ = 52.41, *P* < 0.0001) (Figure 2). The mean numbers of larvae, pupae, adults, and parasitoids were each the highest at 3 weeks and the lowest at 5 weeks after planting. The parasitoids found at site 2 were similar to that found at site 1 and *O. dissitus* was the most abundant among all parasitoids.

### Planting 3 (September-October 2014)

#### Site 1 (bean)

In Planting 3, the mean numbers of mines, larvae, pupae, and adults of *L. trifolii* and the mean numbers of its parasitoids on bean were significantly affected by sample dates (*F*_3, 93_ = 127.58, *P* < 0.0001; Larvae: *F*_3, 93_ = 128.04, *P* < 0.0001; Pupae: *F*_3, 93_ = 122.40, *P* < 0.0001; Adults: *F*_3, 93_ = 102.89, *P* < 0.0001; and Parasitoids: *F*_3, 93_ = 91.09, *P* < 0.0001) (Figure 3). The mean numbers of mines, larvae, pupae, emerged adults, and parasitoids (56.03 ± 4.12, 50.28 ± 4.07, 45.40 ± 3.56, 29.93 ± 2.93, and 9.71 ± 0.89 / 5 leaves; respectively) were all highest at 2 weeks after planting (Sep 20) (Figure 3). Mean numbers of mines, larvae, pupae, and adults of *L. trifolii* and mean numbers of its parasitoids decreased significantly at 3, 4, and 5 weeks after planting with lowest values at 5 weeks after planting (Oct 11) (4.78 ± 0.69; 3.37 ± 0.54; 2.59 ± 0.39; 1.84 ± 0.30; and 0.50 ± 0.12 / 5 leaves; respectively) (Figure 3). The parasitoids recorded at site 1 were *Opius dissitus, Diglyphus sp., Euopius* sp. and *Diaulinopsis callichroma. O. dissitus* was the most abundant parasitoid and was about 75% of total population.

**Figure 3.**
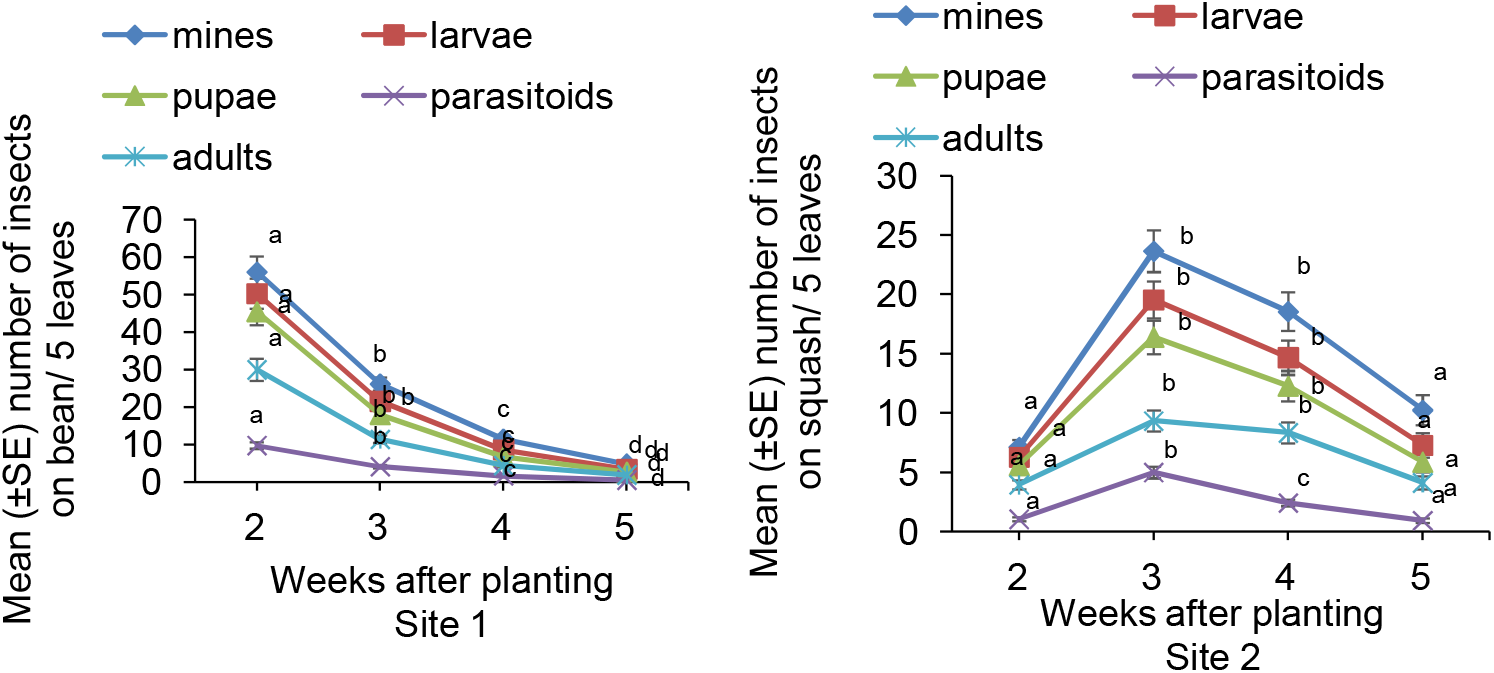
Planting 3 (6 Sep through 11 Oct 2014), Site 1 (bean) and Site 2 (squash) abundance of *L. trifolii* and its parasitoids (mean ± SE / 5 leaves). Means with same letters across the sample dates do not differ significantly based on a Tukey-Kramer test (*P* > 0.05).

#### Site 2 (squash)

Consistent with the two previous plantings, the mean numbers of *L. trifolii* mines, larvae, pupae, adults and parasitoids on squash in Planting 3 were significantly affected by sample dates (Mines: *F*_3, 93_ = 29.59, *P* < 0.0001; Larvae: *F*_3, 93_ = 26.49, *P* < 0.0001; Pupae: *F*_3, 93_ = 23.79, *P* < 0.0001; Adults: *F*_3, 93_ = 16.7, *P* < 0.0001; and Parasitoids: *F*_3, 93_ = 37.27, *P* < 0.0001) (Figure 3).

The mean numbers of mines, larvae, pupae, and adults on squash were significantly higher at 3 and 4 weeks after planting than 2 and 5 weeks after planting (Figure 3). Similarly, mean numbers of parasitoids were significantly higher at 3 weeks after planting (5.00 ± 0.49 parasitoids / 5 leaves) than on other sample dates and were lower (0.93 ± 0.18 parasitoids / 5 leaves) at 5 weeks after planting. *Opius dissitus, Diglyphus sp*., *Euopius* sp. and *Diaulinopsis callichroma* were the parasitoids recorded at site 2. *O. dissitus* was the most abundant parasitoid and was about 70% of total population.

### Planting 4 (November 2014-January 2015)

#### Site 1 (bean)

The mean numbers of *L. trifolii* mines, larvae, pupae, adults and parasitoids on bean were significantly affected by sample dates (Mines: *F*_3, 93_ = 12.04, *P* < 0.0001; Larvae: *F*_3, 93_ = 10.49, *P* < 0.0001; Pupae: *F*_3, 93_ = 6.73, *P* = 0.0005; Adults: *F*_3, 93_ = 3.66, *P* = 0.0165; and Parasitoids: *F*_3, 93_ = 7.71, *P* = 0.0002) (Figure 4). However, the pattern of population density was not consistent and was different from the previous 3 plantings. The mean numbers of mines were significantly higher at 4 weeks after planting (11.37 ± 0.986 mines / 5 leaves, Dec 26) compared with 2 weeks (4.75 ± 0.90 mines / 5 leaves) and 3 weeks after planting (6.79 ± 1.03 mines / 5 leaves) (Figure 4). The mean numbers of larvae, pupae, adults and parasitoids also were significantly higher at 4 weeks compared with 2 weeks and 3 weeks after planting (Figure 4). *Opius dissitus, Diglyphus sp., Euopius* sp. and *Diaulinopsis callichroma. O. dissitus* was the most abundant parasitoid and was about 50% of total population.

**Figure 4.**
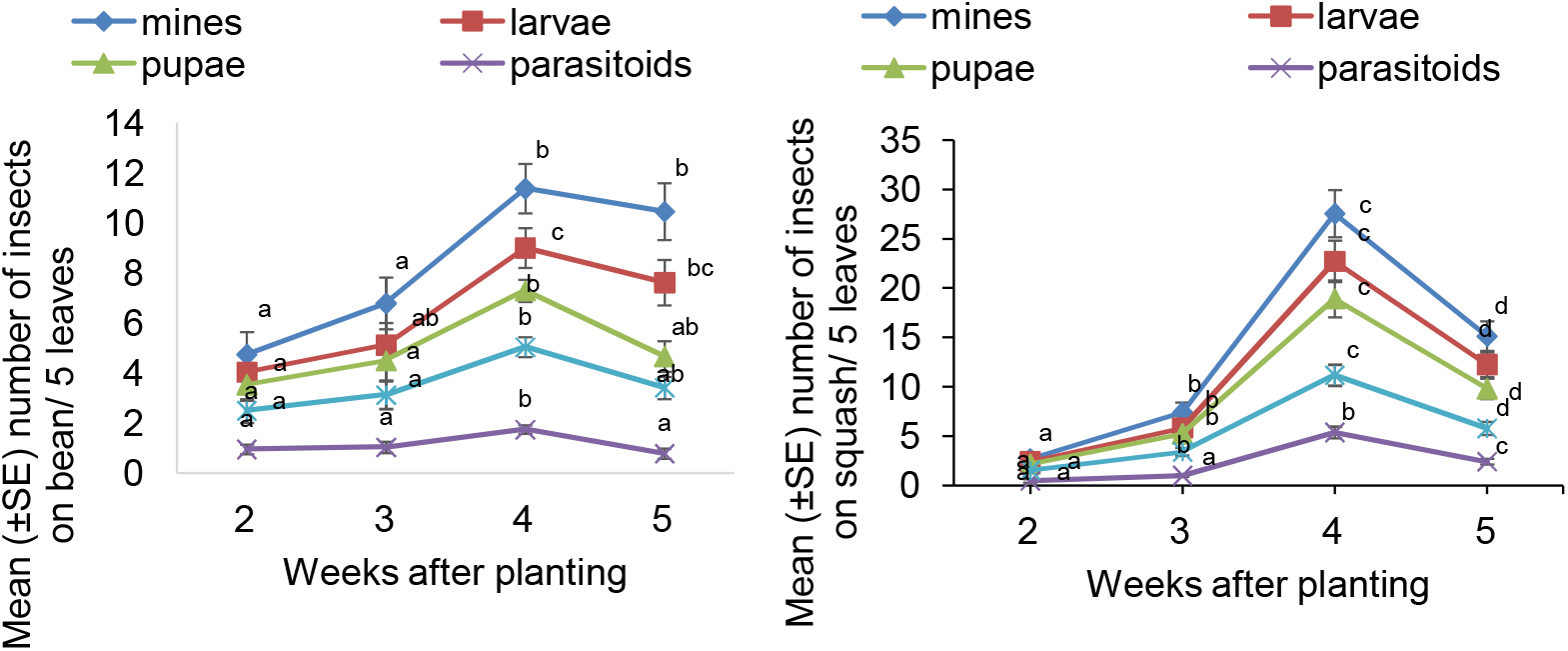
Planting 4 (28 Nov 2014 through 2 Jan 2015), Site 1 (bean) and Site 2 (squash)abundance of *L. trifolii* and its parasitoids (mean ± SE / 5 leaves). Means with same letters across the sample dates do not differ significantly based on a Tukey-Kramer test (*P* > 0.05).

#### Site 2 (squash)

Unlike the other 3 plantings, Planting 4 had different population density trends (Figure 4-8). The mean numbers of mines, larvae, pupae, adults, and parasitoids on squash were significantly affected by sample dates (Mines: *F*_3, 141_ = 69.66, *P* < 0.0001; Larvae: *F*_3, 141_ = 68.76, *P* < 0.0001; Pupae: *F*_3, 141_ = 57.84, *P* < 0.0001; Adults: *F*_3, 141_ = 48.80, *P* < 0.0001; and parasitoids: *F*_3, 141_ = 54.22, *P* < 0.0001) (Figure 4). At 2 weeks after planting the squash (Dec 12), the mean numbers of mines (2.70 ± 0.44 mines / 5 leaves) were significantly lower than the three other sample dates and was the lowest of the planting. However, the mean numbers of mines at 4 weeks after planting were significantly higher than on the other dates (27.54 ± 2.39 mines / 5 leaves) (Figure 4). Similar results were observed for mean numbers of larvae, pupae, adults, and parasitoids with minima and maxima at 2 weeks and 4 weeks after planting, respectively (Figure 4). *O. dissitus* was the most abundant parasitoid and was about 50% of total population of parasitoids.

## Spatial Distribution

### Planting 1 (October - November 2013)

#### Site 1 (bean)

Based on the numbers of mines, leafminer distributions on bean were uniform in all plot sizes (23, 46, and 92 m^2^) at 2 weeks after planting. Slopes *b* and *β* from Taylor’s power law and Iwao’s patchiness regression models were significantly < 1 (Table 1, *P* < 0.05). However, the distribution changed to aggregated for all plot sizes at 3 and 4 weeks after planting. Slopes *b* and *β* were significantly >1 (Table 1, *P* < 0.05). But at 5 weeks after planting, the distribution pattern of mines in the smallest plots (23 m^2^) had mixed results among models with Taylor’s power law results not consistent with Iwao’s patchiness results. However Iwao’s provided better fit of data because of the higher r^2^. Hence, the overall mine distribution pattern for 23 m^2^ plots was apparently aggregated.

**Table 1.** Taylor’s power law and Iwao’s patchiness regression parameters for distribution of *L. trifolii* mines on beans and squash sampled in Planting 1 (Oct – Nov 2013)

| Sample date | Plot size | Bean |  |  |  | Squash |  |  |  |
| --- | --- | --- | --- | --- | --- | --- | --- | --- | --- |
|  |  | Taylor's power law |  | Iwao's patchiness regression |  | Taylor's power law |  | Iwao's patchiness regression |  |
| | | $r^2$ | $b$ | $r^2$ | $\beta$ | $r^2$ | $b$ | $r^2$ | $\beta$ |
| Nov 9 | 23 | 0.001 | -0.004 UNI | 0.835 | 0.883 UNI | 0.001 | -0.126 UNI | 0.006 | 0.796 UNI |
|  | 46 | 0.005 | 0.367 UNI | 0.934 | 0.928 UNI | 0.211 | 0.795 UNI | 0.005 | 0.942 UNI |
|  | 92 | 0.008 | 0.007 UNI | 0.959 | 0.927 UNI | 0.342 | -0.729 UNI | 0.006 | 0.361 UNI |
| Nov 16 | 23 | 0.383 | 4.718 AGG | 0.849 | 1.238 AGG | 0.095 | -2.799 UNI | 0.532 | 0.681 UNI |
|  | 46 | 0.148 | 2.301 AGG | 0.863 | 1.141 AGG | 0.001 | 0.187 UNI | 0.656 | 0.902 UNI |
|  | 92 | 0.446 | 2.691 AGG | 0.917 | 1.196 AGG | 0.382 | 1.288 AGG | 0.964 | 1.027 AGG |
| Nov 23 | 23 | 0.152 | 2.215 AGG | 0.708 | 1.227 AGG | 0.007 | 0.525 UNI | 0.772 | 1.062 AGG |
|  | 46 | 0.081 | 1.817 AGG | 0.541 | 1.041 AGG | 0.201 | 2.174 AGG | 0.839 | 1.195 AGG |
|  | 92 | 0.263 | 2.122 AGG | 0.882 | 1.222 AGG | 1.639 | 1.639 AGG | 0.493 | 1.036 AGG |
| Nov 30 | 23 | 0.013 | 0.781 UNI | 0.653 | 1.150 AGG | 0.007 | 0.265 UNI | 0.220 | 0.800 UNI |
|  | 46 | 0.104 | 1.296 AGG | 0.862 | 1.050 AGG | 0.009 | 0.263 UNI | 0.438 | 0.826 UNI |
|  | 92 | 0.288 | 2.266 AGG | 0.914 | 1.108 AGG | 0.485 | -3.306 UNI | 0.041 | -0.318 UNI |

AGG, aggregated distribution ( $b$ and $\beta$ significantly $>1$ , $P \leq 0.05$ ); RAN, random distribution ( $b$ and $\beta$ not significantly different from 1, $P > 0.05$ ); UNI, uniform distribution ( $b$ and $\beta$ significantly $< 1$ , $P \leq 0.05$ ).

The distribution of parasitoids, however, did not show consistent results. At 2 weeks after planting (Nov 9), both the Taylor’s power law and Iwao’s patchiness regression models contradicted with each other in 23 and 46 m^2^ plots. Because Iwao’s patchiness regression model had a higher r^2^ value and provided a better fit to the data and exhibited random and aggregated distribution, respectively. But for the largest plots, both the linear regression models yielded aggregated distributions (Table 2). At 3 weeks after planting (Nov 16), both linear regression models showed aggregated distributions for 23 and 46 m^2^ plots, but were uniform for large (92 m^2^) plots. However at 4 weeks after planting (Nov 23), the parasitoid distributions in all plot sizes were uniform based on both models (Table 2). At 5 weeks after planting, however (Nov 30), both models contradicted each other for all plot sizes. Because Taylor’s power law had higher r^2^ values for each plot size, it provided better fits to the data, and the overall parasitoid distributions were aggregated, uniform, and random for plot sizes 23, 46, and 92 m^2^, respectively (Table 2).

**Table 2.**
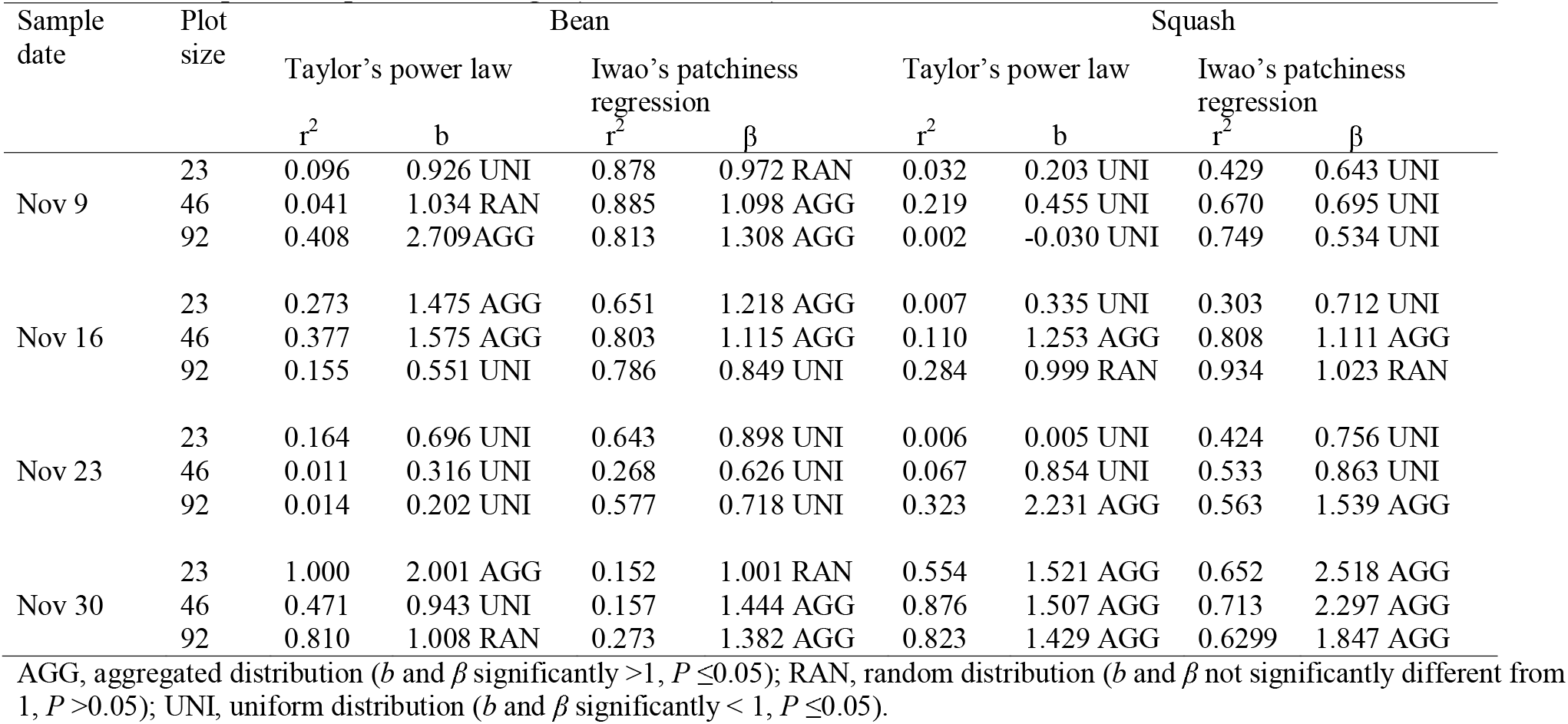
Taylor’s power law and Iwao’s patchiness regression parameters for distribution of parasitoids of *L. trifolii* on bean and squash sampled in Planting 1 (Oct – Nov 2013)

#### Site 2 (squash)

Both the Taylor’s power law and Iwao’s patchiness regression models agreed at 2 weeks after planting and exhibited uniform distributions for leafminers based on numbers of mines in all plot sizes (23, 46 and 92 m^2^) (Table 1). At 3 weeks after planting, the distributions remained uniform in 23 and 46 m^2^ plot, but the distribution for largest plots (92 m^2^) was aggregated. The slopes *b* and *β* were significantly >1 (Table 1, *P* < 0.05). Distributions remained aggregated in all plot sizes at 4 weeks after planting, but changed to uniform at 5 weeks (Table 1). Although both the regression models applied to parasitoid distribution were generally in agreement, there were some inconsistencies (Table 2). The distribution was mostly uniform for 2, 3, and 4 weeks after planting, but was aggregated at 5 weeks.

### Planting 2 (May-June 2014)

#### Site 1 (bean)

Based on the mean numbers of mines, leafminer distributions on bean were dissimilar among regression models at 2 weeks after planting (May 24) (Table 3). Iwao’s patchiness regression model provided higher r^2^ values and better fits to the data, which therefore exhibited uniform, random, and aggregated distributions for 23, 46, and 92 m^2^ plots, respectively. Similarly, at 3 weeks after planting (May 31), values of the indices contradicted each other for distributions in 46 and 92 m^2^ plots. Here, Iwao’s patchiness regression yielded higher r^2^ values and provided better fits to the data, which exhibited random distributions. However for the smallest plots (23 m^2^), both the indices showed aggregated distributions (Table 3). At 4 weeks after planting (June 7), based on Iwao’s patchiness regression model, distributions were random, random, and aggregated for plot sizes of 23, 46 and 92 m^2^, respectively. At 5 weeks after planting (June 14), both modes yielded aggregated distributions for all plot sizes.

**Table 3.** Taylor’s power law and Iwao’s patchiness regression parameters for distribution of *L. trifolii* mines on beans and squash sampled in Planting 2 (May – June 2014)

| Sample date | Plot size | Bean |  |  |  | Squash |  |  |  |
| --- | --- | --- | --- | --- | --- | --- | --- | --- | --- |
|  |  | Taylor's power law |  | Iwao's patchiness regression |  | Taylor's power law |  | Iwao's patchiness regression |  |
| | | $r^2$ | $b$ | $r^2$ | $\beta$ | $r^2$ | $b$ | $r^2$ | $\beta$ |
| Nov 9 | 23 | 0.011 | -0.504 UNI | 0.925 | 0.929 UNI | 0.005 | 0.202 UNI | 0.316 | 0.752 UNI |
|  | 46 | 0.081 | 0.704 UNI | 0.971 | 1.001 RAN | 0.229 | 1.295 AGG | 0.628 | 1.296 AGG |
|  | 92 | 0.250 | 1.261 AGG | 0.973 | 1.041 AGG | 0.489 | 1.567 AGG | 0.644 | 1.159 AGG |
| Nov 16 | 23 | 0.270 | 2.225 AGG | 0.943 | 1.056 AGG | 0.371 | 2.621 AGG | 0.663 | 1.032 AGG |
|  | 46 | 0.266 | 1.722 AGG | 0.909 | 1.018 RAN | 0.360 | 2.105 AGG | 0.843 | 1.337 AGG |
|  | 92 | 0.285 | 1.282 AGG | 0.950 | 0.996 RAN | 0.192 | 1.150 AGG | 0.627 | 1.031 RAN |
| Nov 23 | 23 | 0.350 | 1.243 AGG | 0.913 | 1.023 RAN | 0.368 | 2.893 AGG | 0.760 | 1.281 AGG |
|  | 46 | 0.412 | 0.818 UNI | 0.960 | 0.971 RAN | 0.005 | 0.212 UNI | 0.685 | 0.893 UNI |
|  | 92 | 0.840 | 1.558 AGG | 0.965 | 1.146 AGG | 0.056 | 0.566 UNI | 0.813 | 0.923 UNI |
| Nov 30 | 23 | 0.255 | 1.663 AGG | 0.634 | 1.047 AGG | 0.123 | 1.050 AGG | 0.370 | 1.284 AGG |
|  | 46 | 0.388 | 1.324 AGG | 0.872 | 1.122 AGG | 0.341 | 0.933 UNI | 0.572 | 1.112 AGG |
|  | 92 | 0.442 | 2.118 AGG | 0.468 | 1.234 AGG | 0.859 | 1.461 AGG | 0.943 | 1.468 AGG |

AGG, aggregated distribution ( $b$ and $\beta$ significantly $>1$ , $P \leq 0.05$ ); RAN, random distribution ( $b$ and $\beta$ not significantly different from 1, $P > 0.05$ ); UNI, uniform distribution ( $b$ and $\beta$ significantly $< 1$ , $P \leq 0.05$ ).

For distribution of parasitoids, both indices agreed for all times and plot sizes (Table 4). At 2 weeks after planting, parasitoids showed aggregated, uniform, and uniform distribution for 23, 46, and 92 m^2^ plots, respectively. However, at 3 and at 4 weeks after planting, parasitoids yielded uniform, uniform, and aggregated distributions in 23, 46, and 92 m^2^ plots, respectively. At five weeks after planting, distributions were aggregated for all plot sizes (Table 4).

**Table 4.**
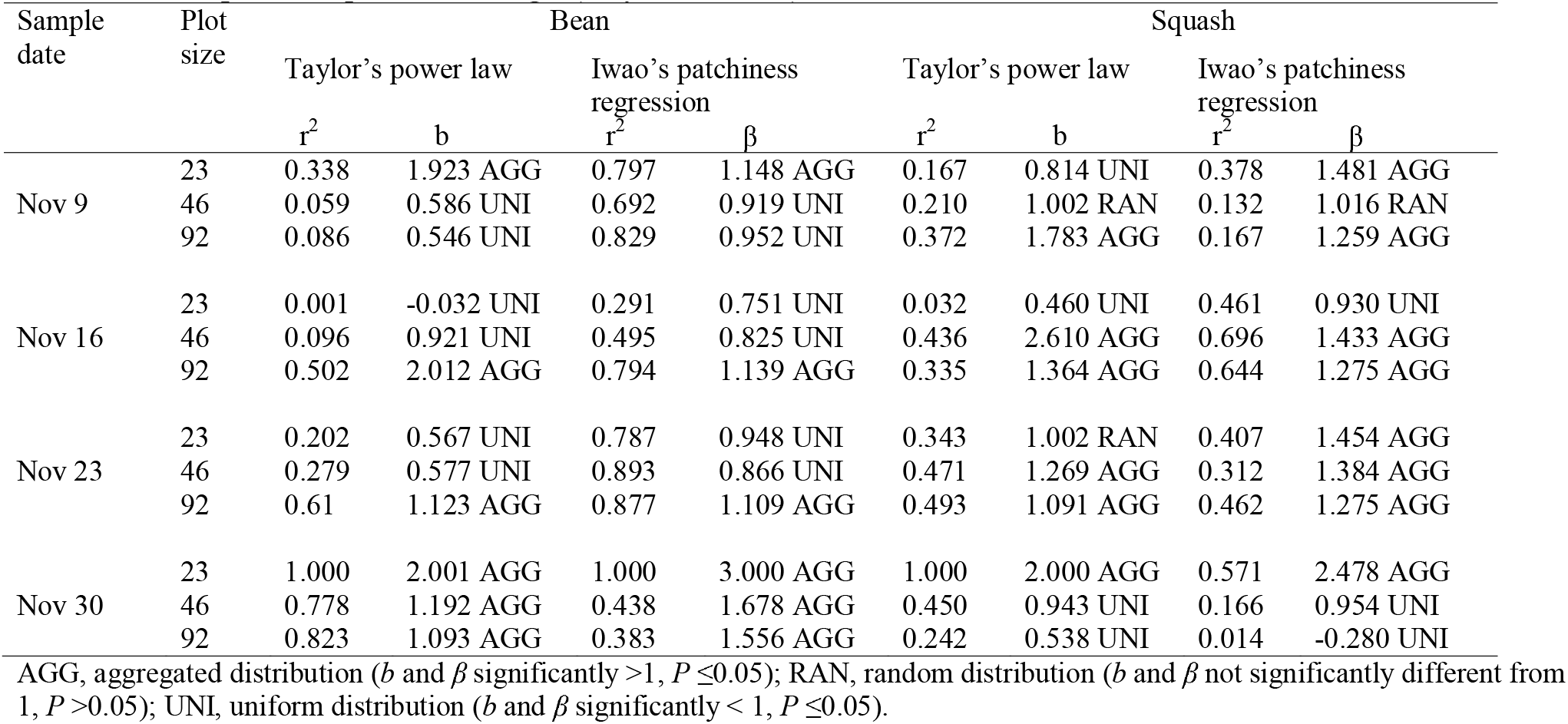
Taylor’s power law and Iwao’s patchiness regression parameters for distribution of parasitoids of *L. trifolii* on bean and squash sampled in Planting 2 (May – June 2014)

#### Site 2 (squash)

Leafminer distributions on squash were not consistent, and the regression models often disagreed with each other (Table 4-3). Because Iwao’s patchiness regression model yielded higher r^2^ values and provided a better fits to the data, I chose its resulting distribution patterns. Distributions of mines at 2 weeks after planting in 23, 46 and 92 m^2^ plots were uniform, aggregated, and aggregated.

But at 3 weeks after planting, the distributions changed to aggregated, aggregated, and random in 23, 46, and 92 m^2^ plots, respectively. However, at 4 weeks after planting, distribution of mines were aggregated, uniform, and uniform in 23, 46, and 92 m^2^ plots, respectively. At 5 weeks after planting, however, distributions of mines were aggregated for all plot sizes.

Parasitoids distributions were mostly aggregated for all plot sizes and sample dates (Table 4). However, at 5 weeks after planting, parasitoid distributions were aggregated in 23 m^2^ plot sizes and uniform in 46 and 92 m^2^ plot sizes.

### Planting 3 (September-October 2014)

#### Site 1 (bean)

Distributions of leafminer mines were not consistent throughout Planting 3 and were contradicting among the regression models at 2 weeks after planting (Sep 20) (Table 5). The plot sizes 23, 46, and 92 m^2^ exhibited uniform, uniform, and aggregated distributions, respectively, because Iwao’s patchiness regression models yielded higher r^2^ values, thus provided better fits to these data. However, at 3 weeks after planting (Sep 27), the models found aggregated distributions for all plot sizes (Table 5). Similarly, at 4 weeks after planting (Oct 4), both indices suggested the distributions were uniform, uniform, and aggregated for 23, 46, and 92 m^2^ plots (Table 5). At 5 weeks after planting (Oct 11), leafminer distributions for 23, 46and 92 m^2^ plots were aggregated, random, and uniform, respectively, based on Iwao’s patchiness regression.

**Table 5.** Taylor’s power law and Iwao’s patchiness regression parameters for distribution of *L. trifolii* mines on beans and squash sampled in Planting 3 (Sep – Oct 2014)

| Sample date | Plot size | Bean |  |  |  | Squash |  |  |  |
| --- | --- | --- | --- | --- | --- | --- | --- | --- | --- |
|  |  | Taylor's power law |  | Iwao's patchiness regression |  | Taylor's power law |  | Iwao's patchiness regression |  |
| | | $r^2$ | $b$ | $r^2$ | $\beta$ | $r^2$ | $b$ | $r^2$ | $\beta$ |
| Sep 20 | 23 | 0.078 | 1.424 AGG | 0.867 | 0.970 UNI | 0.075 | 0.684 UNI | 0.272 | 0.874 UNI |
|  | 46 | 0.001 | -0.089 UNI | 0.920 | 0.879 UNI | 0.107 | -1.272 UNI | 0.061 | 0.301 UNI |
|  | 92 | 0.234 | 1.889 AGG | 0.772 | 1.109 AGG | 0.250 | -2.093 UNI | 0.006 | -0.072 UNI |
| Sep 27 | 23 | 0.135 | 2.752 AGG | 0.882 | 1.315 AGG | 0.532 | 2.585 AGG | 0.746 | 1.406 AGG |
|  | 46 | 0.226 | 4.816 AGG | 0.850 | 1.265 AGG | 0.371 | 1.722 AGG | 0.889 | 1.149 AGG |
|  | 92 | 0.199 | 1.264 AGG | 0.903 | 1.052 AGG | 0.443 | 1.027 RAN | 0.957 | 1.003 RAN |
| Oct 4 | 23 | 0.024 | 0.494 UNI | 0.418 | 0.808 UNI | 0.067 | 1.917 AGG | 0.528 | 1.456 AGG |
|  | 46 | 0.288 | 0.893 UNI | 0.727 | 0.942 UNI | 0.271 | 1.929 AGG | 0.792 | 1.243 AGG |
|  | 92 | 0.699 | 1.347 AGG | 0.928 | 1.095 AGG | 0.284 | -7.203 UNI | 0.049 | -0.736 UNI |
| Oct 11 | 23 | 0.430 | 1.910 AGG | 0.438 | 1.232 AGG | 0.056 | 0.947 UNI | 0.286 | 0.958 UNI |
|  | 46 | 0.164 | 1.226 AGG | 0.388 | 0.990 RAN | 0.007 | -0.391 UNI | 0.059 | 0.606 UNI |
|  | 92 | 0.028 | -0.762 UNI | 0.003 | -0.182 UNI | 0.636 | 1.820 AGG | 0.820 | 1.479 AGG |

AGG, aggregated distribution ( $b$ and $\beta$ significantly $>1$ , $P \leq 0.05$ ); RAN, random distribution ( $b$ and $\beta$ not significantly different from 1, $P > 0.05$ ); UNI, uniform distribution ( $b$ and $\beta$ significantly $< 1$ , $P \leq 0.05$ ).

For parasitoid distributions, both indices were in agreement 2 weeks after planting with aggregated distributions for all plot sizes (Table 6). At 3 weeks after planting, however, parasitoid distributions disagreed among models. Parasitoids had aggregated, aggregated, and uniform distributions in 23, 46, and 92 m^2^ plots, respectively based on Iwao’s patchiness regression model which yielded higher r^2^ values, thus provided better fits to these data. Similarly, at 4 weeks after planting, based on Iwao’s patchiness regression, parasitoids had aggregated, uniform, and uniform distributions in 23, 46, and 92 m^2^ plots, respectively However, at 5 weeks after planting, distributions were uniform for all plot sizes (Table 6).

**Table 6.**
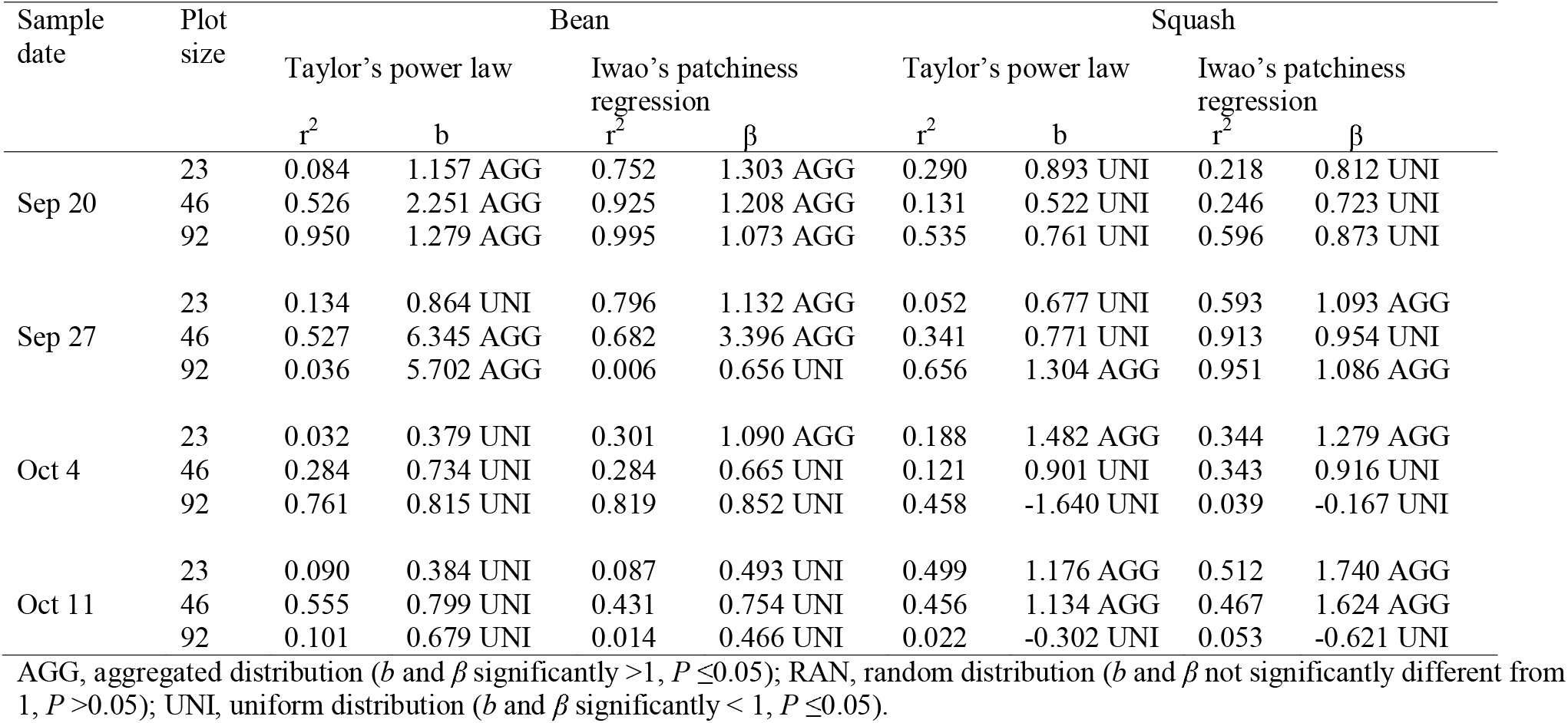
Taylor’s power law and Iwao’s patchiness regression parameters for distribution of parasitoids of *L. trifolii* on bean and squash sampled in Planting 3 (Sep – Oct 2014)

| Sample date | Plot size | Bean |  |  |  | Squash |  |  |  |
| --- | --- | --- | --- | --- | --- | --- | --- | --- | --- |
|  |  | Taylor's power law |  | Iwao's patchiness regression |  | Taylor's power law |  | Iwao's patchiness regression |  |
| | | $r^2$ | $b$ | $r^2$ | $\beta$ | $r^2$ | $b$ | $r^2$ | $\beta$ |
| Sep 20 | 23 | 0.084 | 1.157 AGG | 0.752 | 1.303 AGG | 0.290 | 0.893 UNI | 0.218 | 0.812 UNI |
|  | 46 | 0.526 | 2.251 AGG | 0.925 | 1.208 AGG | 0.131 | 0.522 UNI | 0.246 | 0.723 UNI |
|  | 92 | 0.950 | 1.279 AGG | 0.995 | 1.073 AGG | 0.535 | 0.761 UNI | 0.596 | 0.873 UNI |
| Sep 27 | 23 | 0.134 | 0.864 UNI | 0.796 | 1.132 AGG | 0.052 | 0.677 UNI | 0.593 | 1.093 AGG |
|  | 46 | 0.527 | 6.345 AGG | 0.682 | 3.396 AGG | 0.341 | 0.771 UNI | 0.913 | 0.954 UNI |
|  | 92 | 0.036 | 5.702 AGG | 0.006 | 0.656 UNI | 0.656 | 1.304 AGG | 0.951 | 1.086 AGG |
| Oct 4 | 23 | 0.032 | 0.379 UNI | 0.301 | 1.090 AGG | 0.188 | 1.482 AGG | 0.344 | 1.279 AGG |
|  | 46 | 0.284 | 0.734 UNI | 0.284 | 0.665 UNI | 0.121 | 0.901 UNI | 0.343 | 0.916 UNI |
|  | 92 | 0.761 | 0.815 UNI | 0.819 | 0.852 UNI | 0.458 | -1.640 UNI | 0.039 | -0.167 UNI |
| Oct 11 | 23 | 0.090 | 0.384 UNI | 0.087 | 0.493 UNI | 0.499 | 1.176 AGG | 0.512 | 1.740 AGG |
|  | 46 | 0.555 | 0.799 UNI | 0.431 | 0.754 UNI | 0.456 | 1.134 AGG | 0.467 | 1.624 AGG |
|  | 92 | 0.101 | 0.679 UNI | 0.014 | 0.466 UNI | 0.022 | -0.302 UNI | 0.053 | -0.621 UNI |

AGG, aggregated distribution ( $b$ and $\beta$ significantly $>1$ , $P \leq 0.05$ ); RAN, random distribution ( $b$ and $\beta$ not significantly different from 1, $P > 0.05$ ); UNI, uniform distribution ( $b$ and $\beta$ significantly $< 1$ , $P \leq 0.05$ ).

#### Site 2 (squash)

Both regression models agreed and yielded uniform distributions of leafminer mines in squash at 2 weeks after planting for all plot sizes (Table 5). But 3 weeks after planting, distributions changed to aggregated, aggregated, and random in 23, 46, and 92 m^2^ plots, respectively. Similarly, distributions of mines were aggregated, aggregated, and uniform in 23, 46, and 92 m^2^ plots, respectively at 4 weeks after planting. Distributions changed to uniform, uniform, and aggregated in 23, 46, and 92 m^2^ plot sizes at 5 weeks after planting (Table 5).

Leafminer parasitoid distributions on squash were uniform at 2 weeks after planting for all plot sizes (Table 6). At 3 weeks, the distributions changed to aggregated, uniform, and aggregated for 23, 46, and 92 m^2^ plots. Based on both regression models, mines exhibited aggregated, uniform, and uniform for 23, 46, and 92plots, respectively at 4 weeks after planting. But, at 5 weeks, distributions were aggregated, aggregated, and uniform for 23, 46, and 92 m^2^ plots, respectively (Table 6). Planting 4 (November 2104-January 2015)

Site 1 (bean): Taylor’s and Iwao’s models for distribution of *L. trifolii* mines were in agreement for all plot sizes and sample dates except for the smallest plots (23 m^2^) at 4 week after planting (Table 7). Here, the distributions were uniform for Taylor’s and random for Iwao’s. Distributions in larger plots (46and 92 m^2^) at 4 weeks were aggregated. At 2 weeks after planting (Dec 12), distributions were uniform for the 23 m^2^ plots based on both models. Distributions on larger plots of 46and 92 m^2^ were aggregated. At 3 weeks after planting, distributions aggregated, uniform, and uniform for 23, 46, and 92 m^2^ plots, respectively. At 5 weeks after planting, distributions in 23 m^2^ plots were random, but were uniform in 46and 92 m^2^ plots (Table 7).

**Table 7.** Taylor’s power law and Iwao’s patchiness regression parameters for distribution of *L. trifolii* mines on beans and squash sampled in Planting 4 (Nov 2014 – Jan 2015)

| Sample date | Plot size | Bean |  |  |  | Squash |  |  |  |
| --- | --- | --- | --- | --- | --- | --- | --- | --- | --- |
|  |  | Taylor's power law |  | Iwao's patchiness regression |  | Taylor's power law |  | Iwao's patchiness regression |  |
| | | $r^2$ | $b$ | $r^2$ | $\beta$ | $r^2$ | $b$ | $r^2$ | $\beta$ |
| Dec 12 | 23 | 0.235 | 0.935 UNI | 0.230 | 0.867 UNI | 0.175 | 0.763 UNI | 0.702 | 0.950 UNI |
|  | 46 | 0.711 | 1.663 AGG | 0.550 | 1.435 AGG | 0.590 | 1.454 AGG | 0.823 | 1.312 AGG |
|  | 92 | 0.743 | 1.321 AGG | 0.757 | 1.367 AGG | 0.848 | 1.688 AGG | 0.900 | 1.520 AGG |
| Dec 19 | 23 | 0.296 | 1.838 AGG | 0.350 | 1.156 AGG | 0.031 | 0.486 UNI | 0.343 | 0.925 UNI |
|  | 46 | 0.024 | -0.255 UNI | 0.018 | 0.172 UNI | 0.200 | 1.089 AGG | 0.712 | 1.253 AGG |
|  | 92 | 0.001 | -0.041 UNI | 0.317 | 0.403 UNI | 0.991 | 2.223 AGG | 0.998 | 1.440 AGG |
| Dec 26 | 23 | 0.039 | -1.296 UNI | 0.547 | 1.037 RAN | 0.033 | -0.728 UNI | 0.825 | 0.833 UNI |
|  | 46 | 0.356 | 3.206 AGG | 0.874 | 1.471 AGG | 0.418 | 2.263 AGG | 0.831 | 1.295 AGG |
|  | 92 | 0.775 | 3.360 AGG | 0.930 | 1.428 AGG | 0.209 | -0.014 UNI | 0.999 | 0.793 UNI |
| Jan 2 | 23 | 0.159 | 1.625 AGG | 0.652 | 1.021 RAN | 0.145 | 2.206 AGG | 0.420 | 0.896 UNI |
|  | 46 | 0.039 | 0.286 UNI | 0.877 | 0.861 UNI | 0.298 | 1.185 AGG | 0.847 | 1.002 RAN |
|  | 92 | 0.129 | 0.865 UNI | 0.740 | 0.906 UNI | 0.641 | 1.491 AGG | 0.947 | 1.140 AGG |

AGG, aggregated distribution ( $b$ and $\beta$ significantly $>1$ , $P \leq 0.05$ ); RAN, random distribution ( $b$ and $\beta$ not significantly different from 1, $P > 0.05$ ); UNI, uniform distribution ( $b$ and $\beta$ significantly $< 1$ , $P \leq 0.05$ ).

Similarly, both the Taylor’s power law and Iwao’s patchiness regression models were in agreement for distributions of parasitoids for all plot sizes and sample dates except for the smallest plots (23 m^2^) at 2 week after planting (Table 8). Here, the distributions were uniform for Taylor’s and aggregated for Iwao’s. Distributions in larger plots (46and 92 m^2^) at 2 weeks were aggregated. At 3 weeks after planting, the parasitoids exhibited aggregated, uniform, and uniform distributions for 23, 46, and 92 m^2^ plots, respectively. However, the distributions were uniform for all plot sizes at 4 weeks after planting which changed to aggregated at 5 weeks after planting.

**Table 8.** Taylor’s power law and Iwao’s patchiness regression parameters for distribution of parasitoids of *L. trifolii* on bean and squash sampled in Planting 4 (Nov 2014 – Jan 2015)

| Sample date | Plot size | Bean |  |  |  | Squash |  |  |  |
| --- | --- | --- | --- | --- | --- | --- | --- | --- | --- |
|  |  | Taylor's power law |  | Iwao's patchiness regression |  | Taylor's power law |  | Iwao's patchiness regression |  |
| | | $r^2$ | $b$ | $r^2$ | $\beta$ | $r^2$ | $b$ | $r^2$ | $\beta$ |
| Dec 12 | 23 | 0.221 | 0.823 UNI | 0.294 | 1.408 AGG | 1.000 | 2.000 AGG | 1.000 | 3.000 AGG |
|  | 46 | 0.654 | 1.763 AGG | 0.690 | 1.775 AGG | 0.728 | 1.500 AGG | 0.544 | 2.222 AGG |
|  | 92 | 0.784 | 2.241 AGG | 0.649 | 1.842 AGG | 0.938 | 0.721 UNI | 0.012 | 0.091 UNI |
| Dec 19 | 23 | 0.809 | 1.525 AGG | 0.257 | 1.104 AGG | 0.016 | 0.162 UNI | 0.043 | 0.283 UNI |
|  | 46 | 0.002 | -0.087 UNI | 0.049 | -0.593 UNI | 0.006 | -0.093 UNI | 0.397 | 0.494 UNI |
|  | 92 | 0.012 | -0.189 UNI | 0.074 | -0.494 UNI | 0.006 | 0.035 UNI | 0.679 | 0.292 UNI |
| Dec 26 | 23 | 0.118 | -0.714 UNI | 0.433 | 0.729 UNI | 0.186 | -1.870 UNI | 0.491 | 0.737 UNI |
|  | 46 | 0.001 | -0.091 UNI | 0.725 | 0.741 UNI | 0.008 | 0.352 UNI | 0.625 | 0.943 UNI |
|  | 92 | 0.059 | -0.549 UNI | 0.689 | 0.720 UNI | 0.388 | 0.288 UNI | 0.987 | 0.787 UNI |
| Jan 2 | 23 | 0.944 | 1.552 AGG | 0.839 | 1.523 AGG | 0.001 | 0.141 UNI | 0.196 | 0.805 UNI |
|  | 46 | 0.784 | 1.528 AGG | 0.698 | 1.544 AGG | 0.111 | 1.060 AGG | 0.642 | 1.117 AGG |
|  | 92 | 0.999 | 2.474 AGG | 0.999 | 3.118 AGG | 0.193 | 1.547 AGG | 0.443 | 1.116 AGG |

AGG, aggregated distribution ( $b$ and $\beta$ significantly $>1$ , $P \leq 0.05$ ); RAN, random distribution ( $b$ and $\beta$ not significantly different from 1, $P > 0.05$ ); UNI, uniform distribution ( $b$ and $\beta$ significantly $< 1$ , $P \leq 0.05$ ).

Site 2 (squash): Distributions of *L. trifolii* based on numbers of mines according to both the regression models were consistent throughout all sample dates for all plot sizes, except for the plots of 23and 46 m^2^ at 5 weeks after planting (Table 7). Since, Iwao’s patchiness regression models yielded higher r^2^ values, thus provided better fits to these data and exhibited uniform and random, respectively. Distribution for 92 m^2^ plots were aggregated.

The distribution of parasitoids was mostly uniform except at 2 and 5 weeks after planting (Table 8). At 2 weeks after planting, the distribution was aggregated, aggregated, and uniform for 23, 46, and 92 m^2^ plots, respectively. At 3 weeks, distributions were uniform. At 4 weeks, distributions were uniform except for 46 m^2^ plots, which were uniform using Taylor’s, but were random using Iwao’s, which had the higher r^2^ suggesting it was the most valid result for 46 m^2^ plots. At 5 weeks, results were uniform for 23 m^2^ plots and aggregate for 92 m^2^ plots. Plots of 46 m^2^ were random using Taylor’s and aggregated using Iwao’s, which had a higher r^2^ suggesting it was the most valid result for 46 m^2^ plots (Table 8).

## Discussion

The mean numbers of mines, larvae, pupae, and adult *L. trifolii* on bean were observed to be highest at 2 weeks after planting for all trials except for Planting 4 (Nov 2014 - Jan 2015) (Figures 1, 2, 3, and 4). Similarly, on squash, mean numbers of mines, larvae, pupae, and adults were highest at 3 weeks after planting for all trials, except for Planting 4 (Nov 2014 - Jan 2015) (Figures 1, 2, 3, and 4). With beans, leafminer activity peaked few days after the plants had two primary leaves fully unfolded, and there was significant decrease in leaf miner activity thereafter. In contrast, leafminer activity on squash gradually increased, tending to peak 3 weeks after planting, then it gradually decreased 4 and 5 weeks after planting. These leafminer activity patterns may have resulted from factors such as leaf nutrients, defensive compounds, trichome presence, and cuticle thickness. Many reports have demonstrated the differential leaf utilization by leafminer based on these leaf characteristics (Stiling et al. 1982, Fagoonee and Toory 1983, Nuessly and Nagata 1994, Li et al. 1997, Scheirs et al. 2001, Facknath, 2005, Digweed 2006, Ayabe and Shibata 2008). Results from our study suggest that leafminers on bean prefer the first pair of leaves, which are cotyledonus. Similarly, Chandler and Gilstrap (1987) working with *L. trifolii* on peppers observed an initial period of increased damage during the cotyledon growth phase. Many other studies confirm the exclusive utilization of new (young) leaves of host plants by different leafminer species (Auerbach & Simberloff 1984, Hespenheide 1991, Ayabe and Shibata 2008).

The average across all plantings for leafminer mines, larvae, pupae, and adults on beans were inconsistent (Figure 5). Planting 1 (Oct - Nov 2013) and Planting 3 (Sep - Oct 2014) had similar seasonal averages for numbers of mines, larvae, pupae, and adults. However, Planting 2 (May - June 2014) had apparently higher seasonal averages for mean numbers of mines, larvae, pupae, and adults than Plantings 1 or 3, whereas Planting 4 (Nov 2014 - Jan 2015) had the lowest. However on squash, seasonal averages for mines, larvae, pupae, and adults were similar (comparable) in Plantings 2, 3 and 4, and slightly higher in Planting 1 than the other plantings (Figure 6). Because there was a large temperature decrease in December 2014 during the 4^th^ planting, this low temperature may have helped to minimize the leafminer infestations (<1 mines per leaf). The temperature at 2 weeks after the fourth planting dropped to 17 °C, which was relatively low compared to other sample dates in the planting, when temperatures were generally between 20 - 26°C (FAWN 2015) (Figure 7).

**Figure 5.**
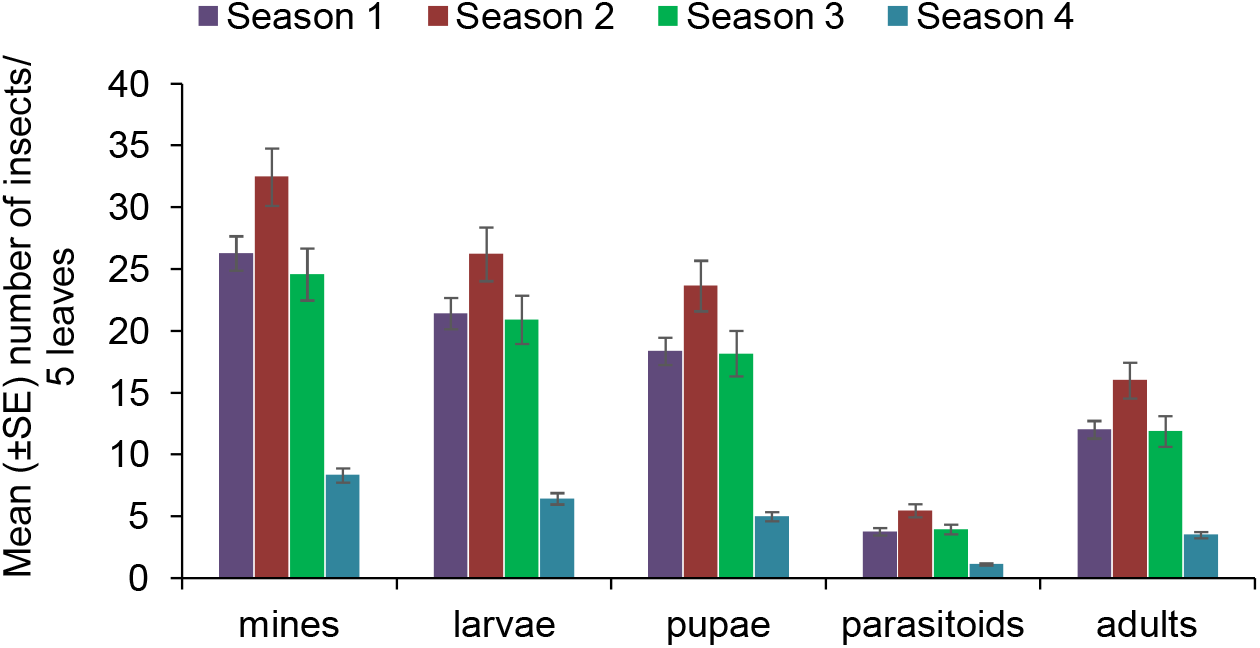
Seasonal abundance (mean ± SE / 5 leaves) of *L. trifolii* mines, larvae, pupae, adults and its parasitoids on bean during 4 plantings.

**Figure 6.**
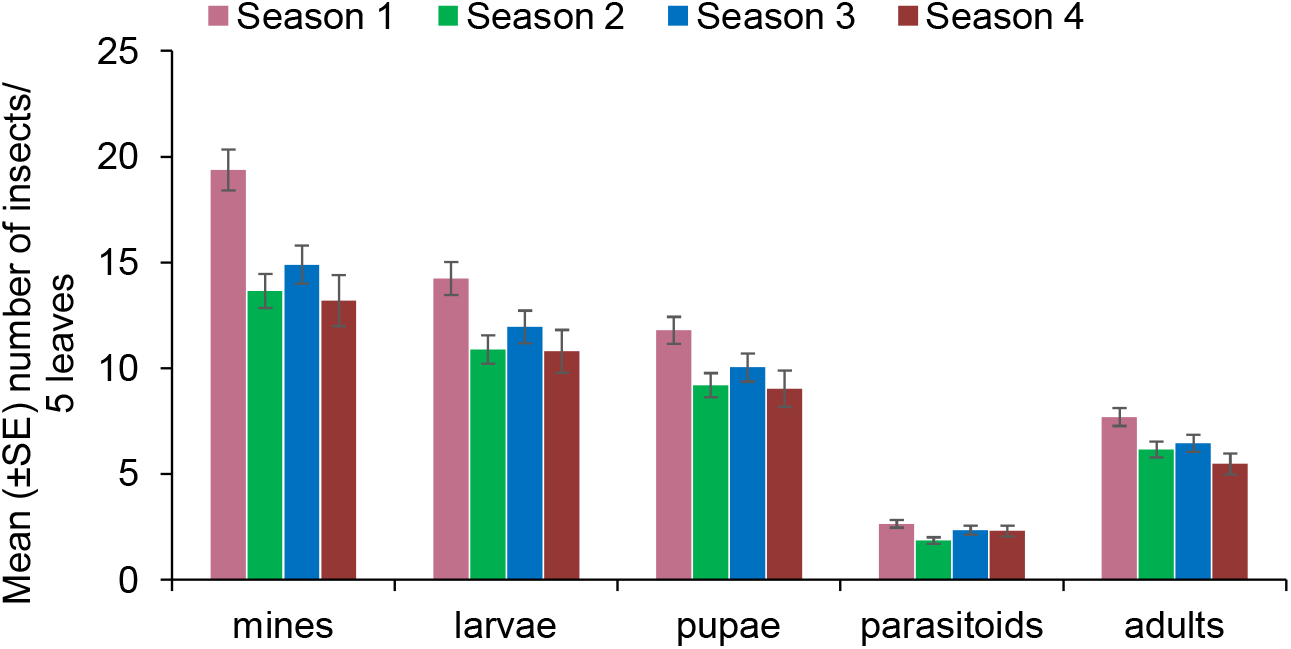
Seasonal abundance (mean ± SE / 5 leaves) of *L. trifolii* mines, larvae, pupae, adults and its parasitoids on squash during 4 plantings.

**Figure 7.**
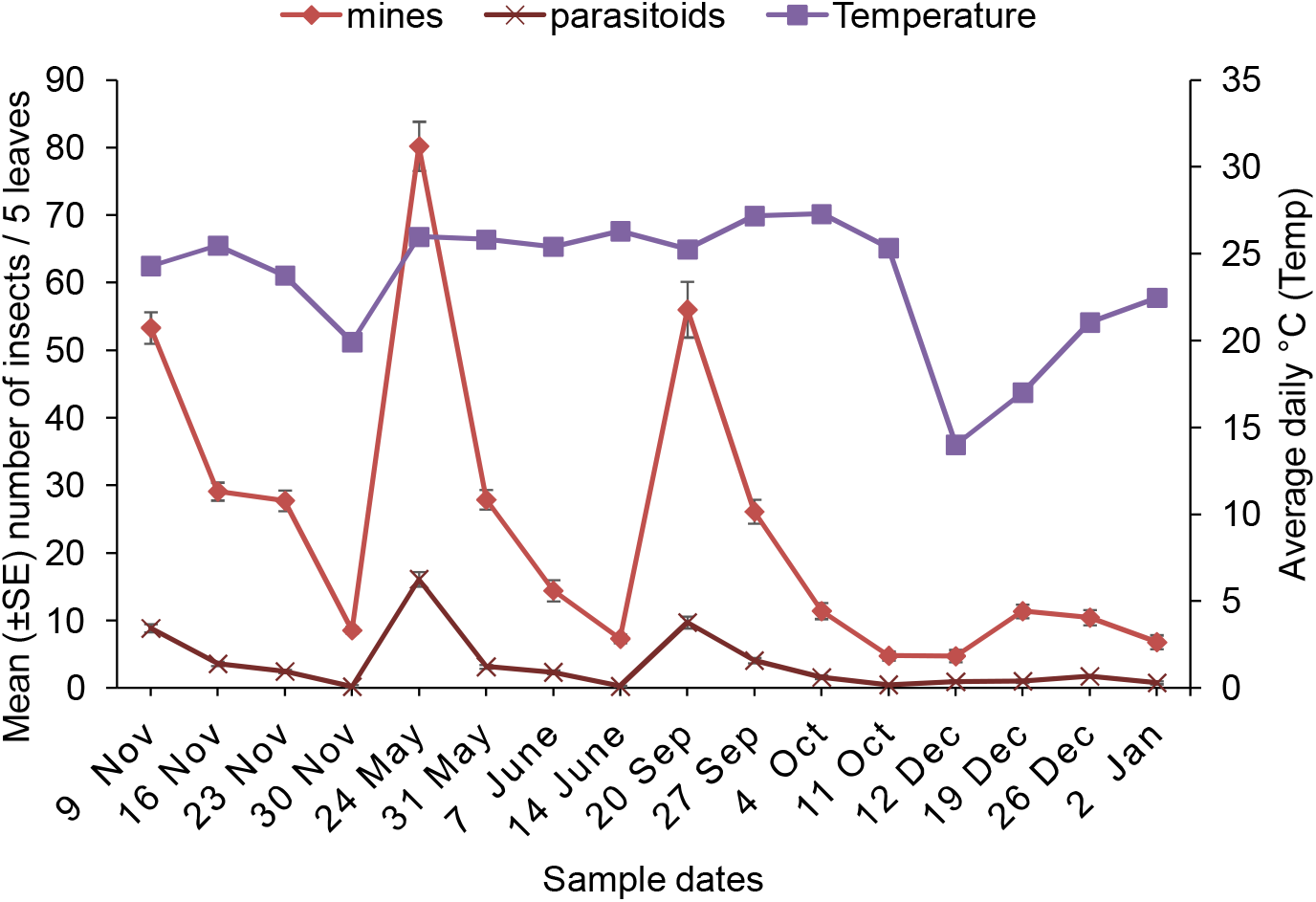
Comparison of average daily temperature (°C) and abundance of *L. trifolii* mines, and parasitoids on bean during the four plantings (26 Oct 2013 – 2 January, 2015).

Similar trends of temperature-dependent fluctuation in population density have been shown in other studies (Johnson et al. 1980, Nestel et al. 1994, Palumbo et al. 1994, Hammad and Nemer 2000, Park et al. 2001, Weintraub 2001, Tran et al. 2005, Arida et al. 2013). Nestel et al. (1994) reported that leafminer populations peaked during intermediate temperatures. They suggested that dynamics of tropical insect populations can be changed with slight variations in climatic conditions in tropical regions. Palumbo et al. (1994) reported decreased leafminer populations in December and January (when temperatures were low) and increased populations in September and October, when temperatures were above 23°C. This pattern of population density was concurrent with our results.

Park et al. (2001) reported a similar temperature-dependent population trend for *L. trifolii* in Korea. They observed that populations of leafminer adults increased immediately after transplanting *Gerbera jamesonii* in April, and the population was still higher in mid-May, early September, and late October while the population decreased in December. Similarly, Tran et al. (2005) reported that leafminer populations in Vietnam were highest in November with densities as high as 38 larvae/leaf. Similarly, Hammad and Nemer (2000) reported that leafminer population densities were lower with temperatures above 28 °C, but were relatively high with temperatures of 20-27 °C. In our study, the leafminer populations were highest within a similar temperature range (24-26 °C).

Population densities of *L. trifolii* parasitoids showed similar trends as leafminers on both bean and squash. Population densities of the parasitoids were highest when leafminer population densities were also high. Occurrence of parasitism and its magnitude varies with leafminer densities (Palumbo et al. 1994). Hence, this parasitism by leafminer parasitoids may be density dependent parasitism, which may warrant further investigation.

Based on numbers of larvae sampled, *L. trifolii* populations on squash and bean did not show a particular distribution pattern on all the sample dates. The distribution patterns appeared similar on both crops. *L. trifolii* exhibited mostly aggregated distributions on each crop on most sample dates. Similar results were reported by Beck et al. (1981) Jones & Parrella (1986), and Hammad and Nemer (2000). Distributions of parasitoids and their leafminer hosts were similar on bean and squash. Therefore, these results do not provide enough evidence to conclude that distribution of parasitoids is affected by weather parameters instead we can conclude that the distribution is depended on leafminer density.

Results of the present study indicated that leafminer preferred certain chronologies of bean and squash plantings over others. Perhaps additional studies should investigate differences in the physical and chemical properties of bean and squash leaves at different periods after planting and any differential effects they may have on leafminer parasitism.

## Acknowledgments

We thank R. Rijal-Devkota, C. M. Sabines, B. Panthi, C. Carter,and J. Teyes for assistance.

